# Deferiprone, an Iron Chelator, is Preventive and Therapeutic in Experimental Crescentic Glomerulonephritis

**DOI:** 10.1101/225540

**Authors:** Tai-Di Chen, Jeong-Hun Ko, Maria Prendecki, Stephen P. McAdoo, Charles D. Pusey, H. Terence Cook, Jacques Behmoaras

## Abstract

Crescentic glomerulonephritis represents the most severe form of antibody-mediated glomerulonephritis. It is an important cause of renal dysfunction worldwide and there is a need for more effective treatment. Deferiprone, an orally active iron chelator, is widely used in patients with thalassemia. Here we present the preventive and therapeutic effects of deferiprone in experimental crescentic glomerulonephritis. Nephrotoxic nephritis was induced in Wistar Kyoto rats, and preventive treatment with deferiprone substantially lowered glomerular crescent formation by 84%, with 70% reduction in proteinuria. In established glomerulonephritis, deferiprone treatment effectively halted glomerular inflammation, reversed progression of proteinuria, and prevented deterioration of renal function. Deferiprone reduced glomerular inflammatory cell proliferation *in vivo.* It was internalised by monocyte/macrophages and inhibited their proliferation *in vitro,* without showing cellular toxicity. Interestingly, deferiprone showed a neutralizing effect on superoxide anions, and prevented the expression of monocyte chemoattractant protein-1 and matrix metalloproteinase 9, 12 and 14, by primary macrophages. These results suggest that deferiprone partly exerts its renal protective effect through inhibition of monocyte/macrophage proliferation and function by iron-chelating and anti-oxidant properties, respectively. We conclude that deferiprone is an effective treatment in a severe and reproducible model of antibody-mediated glomerular inflammation that resembles human crescentic glomerulonephritis, indicating its therapeutic potential.

## Introduction

Crescentic glomerulonephritis is generally characterized by a sudden decline in renal function. Clinically, it is often referred to as rapidly progressive glomerulonephritis, and can be seen in a range of different diseases with diverse aetiologies and underlying pathogeneses.^1,2^ The histologic hallmark of crescentic glomerulonephritis is disruption of the glomerular basement membrane (GBM) with accumulation of proliferating macrophages and parietal epithelial cells in Bowman’s space, forming crescents. The diagnosis of crescentic glomerulonephritis comprises clinical, histological, and serological studies, and irrespective of its aetiology, its prognosis is generally considered unfavourable.^3^ Currently there is no specific or definitive therapy for crescentic glomerulonephritis. Glucocorticoids, cytotoxic agents, and plasmapheresis can be effective, but lead to serious side effects, especially in a prolonged treatment course.^4^

Nephrotoxic nephritis (NTN) in the uniquely susceptible Wistar Kyoto (WKY) rat is a well-established and reproducible model of crescentic glomerulonephritis. It was first described by Granados *et al.,* who showed rapid development of severe crescentic glomerulonephritis in WKY rats following a single injection of nephrotoxic serum raised in rabbits.^5^ This model, which resembles the human pathology, was further characterized by others,^6–8^ and the role of infiltrating monocyte/macrophages was found to be critical in the development of the disease, as shown by monocyte/macrophage depletion and treatment with antibodies against monocyte chemoattractant protein-1 (MCP-1)^9,10^. Our group has contributed to the understanding of the role of macrophages in crescentic glomerulonephritis by using a genetic approach in the WKY rat, and found that the two most significant quantitative trait loci linked to glomerular crescent formation control macrophage function.^11–14^ Recently, in an attempt to genetically map the remaining crescentic glomerulonephritis loci, we identified macrophage ceruloplasmin, a copper-carrying protein, as a positional candidate regulating macrophage activity, possibly through its ferroxidase activity and regulation of intracellular iron. This latter finding suggested that macrophage iron metabolism could contribute to the pathogenesis of crescentic glomerulonephritis.^15^

Iron participates in many aspects of cellular biology and is essential for life as its deprivation leads to inactivation of ribonucleotide reductase (RNR), the enzyme essential for DNA synthesis.^16–18^ Thus iron chelation emerges as an alternative anti-cancer and immunosuppressive therapy.^18,19^ Iron is also an active participant in the generation of superoxide and other reactive oxygen species (ROS),^20^ which have been shown to damage cellular components such as DNA and proteins.^21^ Several studies have shown that oxidative stress plays a crucial role in the development of various renal diseases, ranging from crescentic glomerulonephritis to diabetic nephropathy, both in animal models and in humans.^22–24^ In addition to their direct cell-damaging consequences, superoxide radicals also serve as secondary messengers for downstream cell responses in producing monocyte chemoattractant protein-1 (MCP-1) and matrix metalloproteinases (MMPs),^25,26^ which participate in the development of crescentic glomerulonephritis.^9,27^

Deferiprone is an iron chelating drug widely used for the treatment of iron overload in patients with transfusion-dependent thalassemia.^28^ It is also a potent antioxidant.^29^ In this study, we show that administration of deferiprone before the induction of NTN prevents the development of crescentic glomerulonephritis in WKY rats. We propose that deferiprone exerts its protective effects by both its iron-chelating and anti-oxidative properties. *In vivo,* deferiprone inhibited intraglomerular cell proliferation, prevented monocyte/macrophage accumulation, and down-regulated *Mmp9, Mmp12* and *Mmp14* expression in nephritic glomeruli. *In vitro,* deferiprone reduced monocyte/macrophage proliferative activity, neutralized superoxide anions, and prevented expression of *Ccl2* (MCP-1), *Mmp9, Mmp12* and *Mmp14* in activated bone marrow derived macrophages (BMDMs). Furthermore, treatment with deferiprone in established glomerulonephritis was still effective and reversed the progression of proteinuria, preventing further deterioration of renal function. Taken together, these findings suggest that deferiprone could be used in the management of crescentic glomerulonephritis, which is characterized by enhanced proliferation, recruitment, and activation of macrophages.

## Results

### Deferiprone Prevents the Development of Nephrotoxic Nephritis

To investigate whether deferiprone prevents the development of experimental crescentic glomerulonephritis, WKY rats were treated with either vehicle or deferiprone (200mg/kg/day, starting one hour before the injection of nephrotoxic serum, until day 6 following NTN induction). The disease severity was assessed on day 7. (Figure 1A).

**Figure 1.**
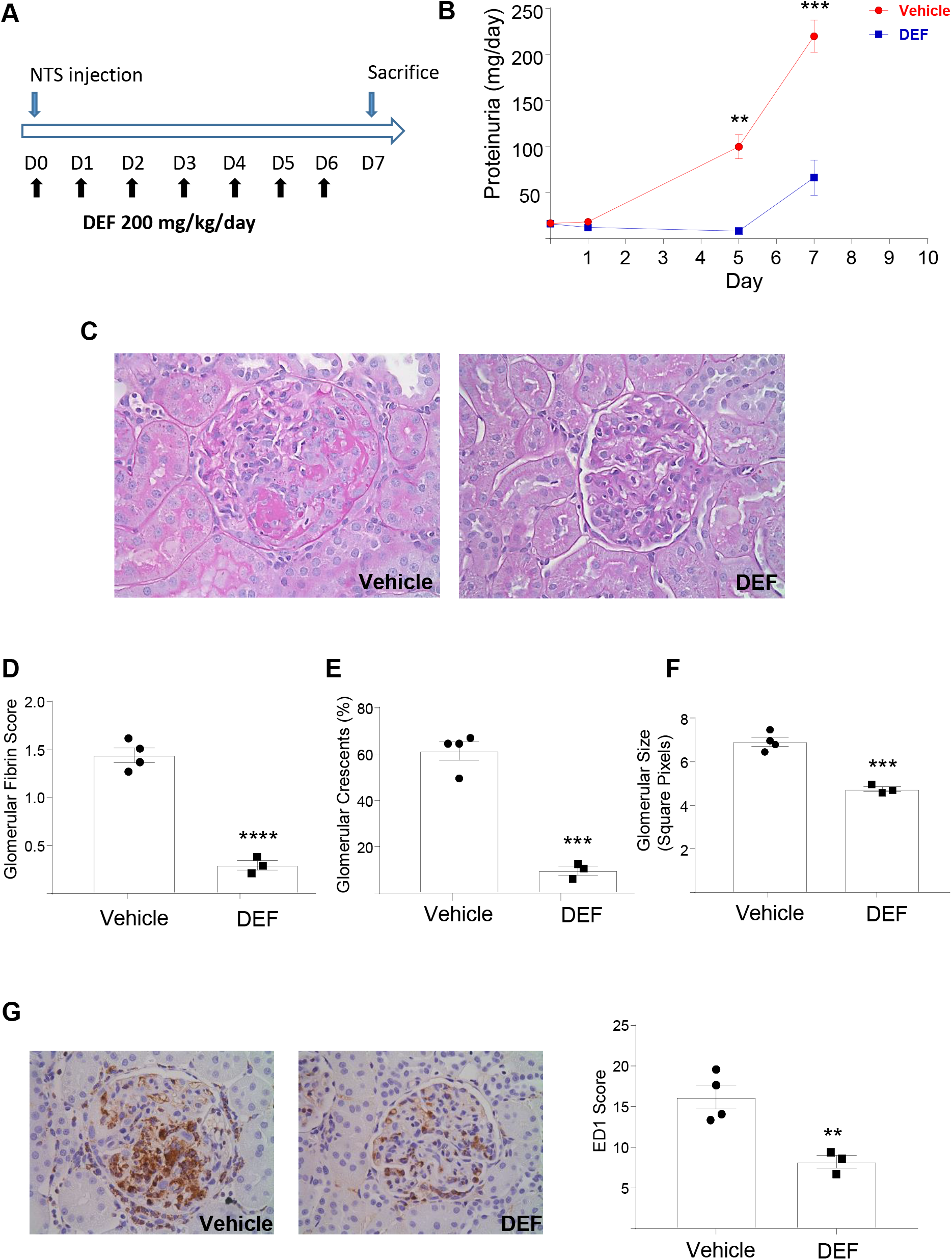
Preventive NTN experiment using deferiprone in the WKY rats. (A) Experimental design. (B) Amount of urinary protein at different time points during the course of NTN. (C) PAS staining showing representative glomerular morphology of the vehicle (left) and the deferiprone (DEF) groups (right). (D) Glomerular fibrin score. (E) Percentage of glomeruli with crescents. (F) Glomerular size. (G) ED1 staining showing macrophages in the glomeruli; ED1 score was normalized to glomerular size. Photographs original magnification, ×400. n=4 for the vehicle and n=3 for the deferiprone (DEF) groups. \**P* ≤ 0.05; ≤\*\**P* ≤ 0.01; \*\*\**P* ≤ 0.001; \*\*\*\**P* ≤ 0.0001.

Baseline urinary protein levels were identical in both groups before the injection of nephrotoxic serum. As previously shown in NTN-susceptible WKY rats, proteinuria rapidly develops in the vehicle-treated group, reaching 220±18 mg/day on day 7 after the induction of NTN. In sharp contrast, rats treated with deferiprone had markedly lower levels of proteinuria during the entire disease course (Figure 1B).

Furthermore, the glomerular morphology in each group at sacrifice was different (Figure 1C). Most glomeruli in vehicle-treated rats were markedly enlarged, containing infiltrating cells, showing disruption of capillary walls with fibrin deposits, and displaying prominent extracapillary hypercellularity with large crescents. In contrast, the glomeruli of rats treated with deferiprone were well preserved, with only a few of them showing fibrin deposits or small crescents. Semi-quantitative evaluation of histologic parameters showed that the glomeruli of deferiprone-treated group had a substantially lower fibrin score (Figure 1D). Similarly, crescent formation was drastically reduced by deferiprone treatment, and glomeruli of the control group were larger than the deferiprone-treated group (Figure 1E and F). Since the WKY NTN model depends on the infiltration and activation of monocyte/macrophages,^10^ we performed ED-1 (rat CD68) staining and showed that deferiprone significantly lowered the number of monocyte/macrophages (Figure 1G), which are the main inflammatory cells observed both in human and experimental crescentic glomerulonephritis.^30^

### Deferiprone Inhibits Intraglomerular Cell Proliferation and Macrophage MCP-1 Production

In glomerulonephritis, the larger glomeruli with increased glomerular cellularity is the combined result of inflammatory cell recruitment and local cell proliferation.^31,32^ Intracellular iron is essential for cell proliferation since it is indispensable for the activity of ribonucleotide reductase (RNR), which controls the rate-limiting step in DNA synthesis.^17^ Deferiprone treatment significantly reduced intraglomerular cell proliferation *in vivo* as shown by proliferating cell nuclear antigen (PCNA) staining (Figure 2A). The iron chelation effect of deferiprone was confirmed by the dose-dependent up-regulation of *Tfrc* (transferrin receptor) mRNA expression in BMDMs (Figure 2B), a method previously used for detecting cellular responses to reduced intracellular iron.^18^ The usage of deferiprone up to 500 μM, which corresponds to the commonly used dosage in humans,^29^ did not affect BMDM viability (Figure 2C). In addition, the anti-proliferative effect of deferiprone was further confirmed in monocytic THP-1 cells *in vitro,* where deferiprone treatment resulted in 66% reduction in cell division (Figure 2D).

**Figure 2.**
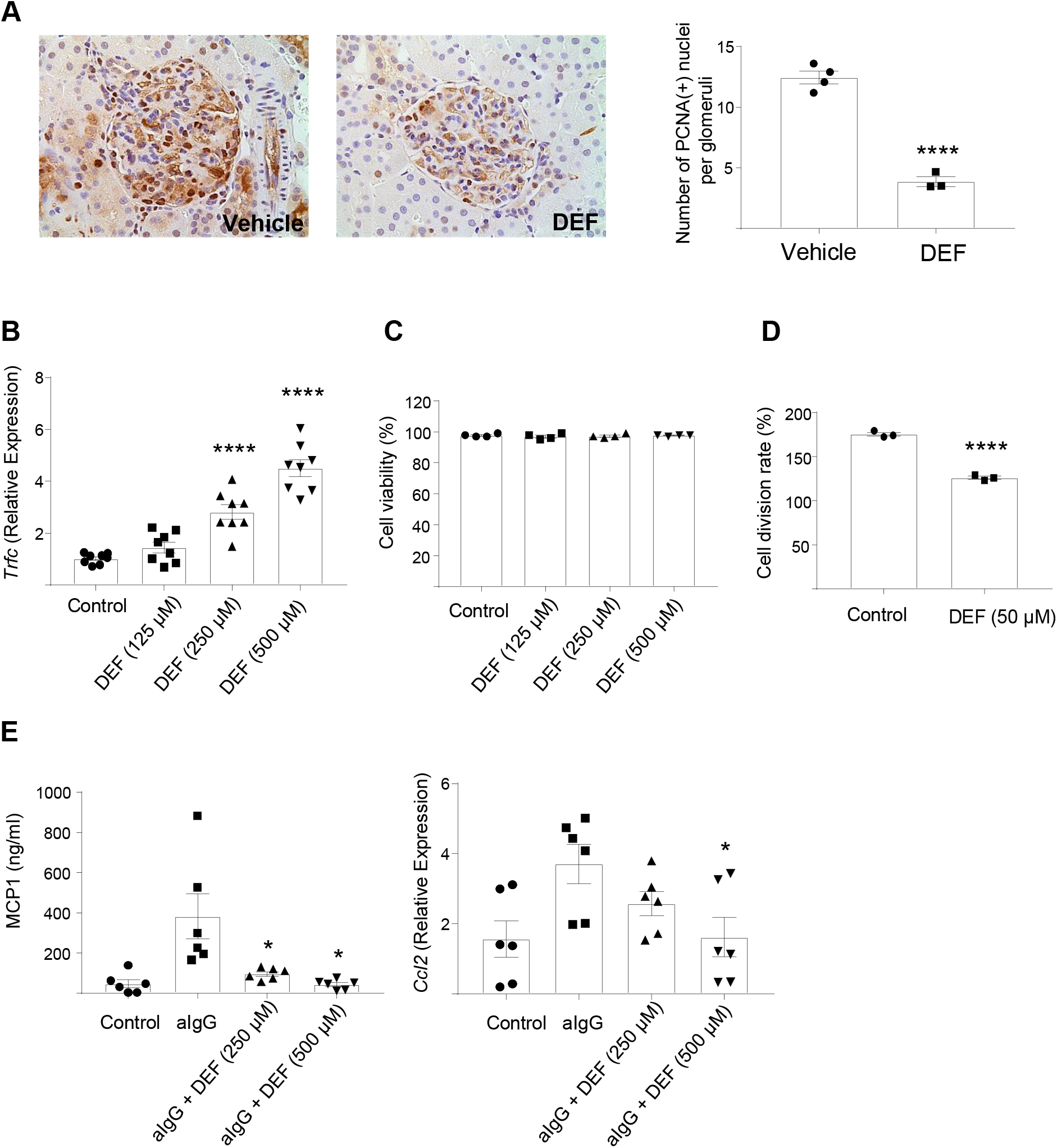
Deferiprone inhibits cell proliferation and MCP-1 production. (A) PCNA staining showing proliferating cells in nephritic glomeruli; number of nuclear PCNA(+) cells in the glomeruli. (B) Transferrin receptor (Tfrc) mRNA expression in BMDMs following 36 hours of incubation with indicated concentrations of deferiprone (DEF). (C) BMDM viability following 36 hours of incubation with different concentrations of deferiprone (DEF). (D) Cell division rate of THP-1 cells in culture medium with or without deferiprone 50 μM. (E) MCP-1 protein levels in cell culture supernatants (left panel) and *Ccl2* mRNA expression (right panel) of BMDMs pre-treated with DEF or control for 2 hours and then stimulated with heat-aggregated IgG (aIgG) for 24 hours. Photographs original magnification, ×400. For all experiments, each group has at least 3 biological replicates with or without technical duplicates. \**P* ≤ 0.05; \*\**P* ≤ 0.01; \*\*\**P* ≤ 0.001; \*\*\*\**P* ≤ 0.0001.

In proliferative glomerulonephritis, macrophages are the most common inflammatory cells found in the glomeruli, and MCP-1 is a crucial chemokine facilitating their recruitment.^9,33,34^ We therefore examined the effect of deferiprone on MCP-1 production in macrophages, which have been previously linked to glomerular injury in nephrotoxic nephritis.^9,10,34^ We found that deferiprone had a dose-dependent effect in reducing MCP-1 protein synthesis and mRNA expression in BMDMs stimulated with heat-aggregated IgG (Figure 2E). Taken together, these results suggest that deferiprone decreased MCP-1 mediated cell recruitment and cell proliferation in NTN, preventing inflammatory cell accumulation and further glomerular damage. Because MCP-1 has been shown to be under the regulation of ROS, we next investigated the effect of DEF on superoxide production in macrophages.^25^

### Deferiprone Ameliorates Oxidative Stress by Directly Neutralizing Superoxide Anions

In the WKY model of nephrotoxic nephritis, macrophage activation by Fc receptor–mediated oxidative burst partly explains the unique susceptibility of this strain to NTN.^12,13^ Superoxide anions are the precursor of most other ROS,^35^ and their production by macrophages has been shown to be a prominent contributor to the glomerular injury in glomerulonephritis.^15,36,37^ To test whether deferiprone protects cells from superoxide induced oxidative stress, we stimulated BMDMs with phorbol 12-myristate 13-acetate (PMA), which is a widely used superoxide inducer in innate immune cells such neutrophils and macrophages.^38,39^ Deferiprone markedly decreased the superoxide anions detected in stimulated BMDMs (Figure 3A). Since deferiprone could decrease the production of superoxide anions by inhibiting its upstream signalling pathway through phosphorylation of protein kinase C α (PKCα),^40^ we next measured PKC phosphorylation in presence of deferiprone but did not find any effect on PKCα activation (Figure 3B). In contrast, deferiprone nearly completely abolished the superoxide anions produced by xanthine-xanthine oxidase reaction in a cell free setup (Figure 3C), suggesting that it could neutralize superoxide anions, irrespective of their source, and prevent downstream oxidative stress-dependent MCP-1 production in macrophages.

**Figure 3.**
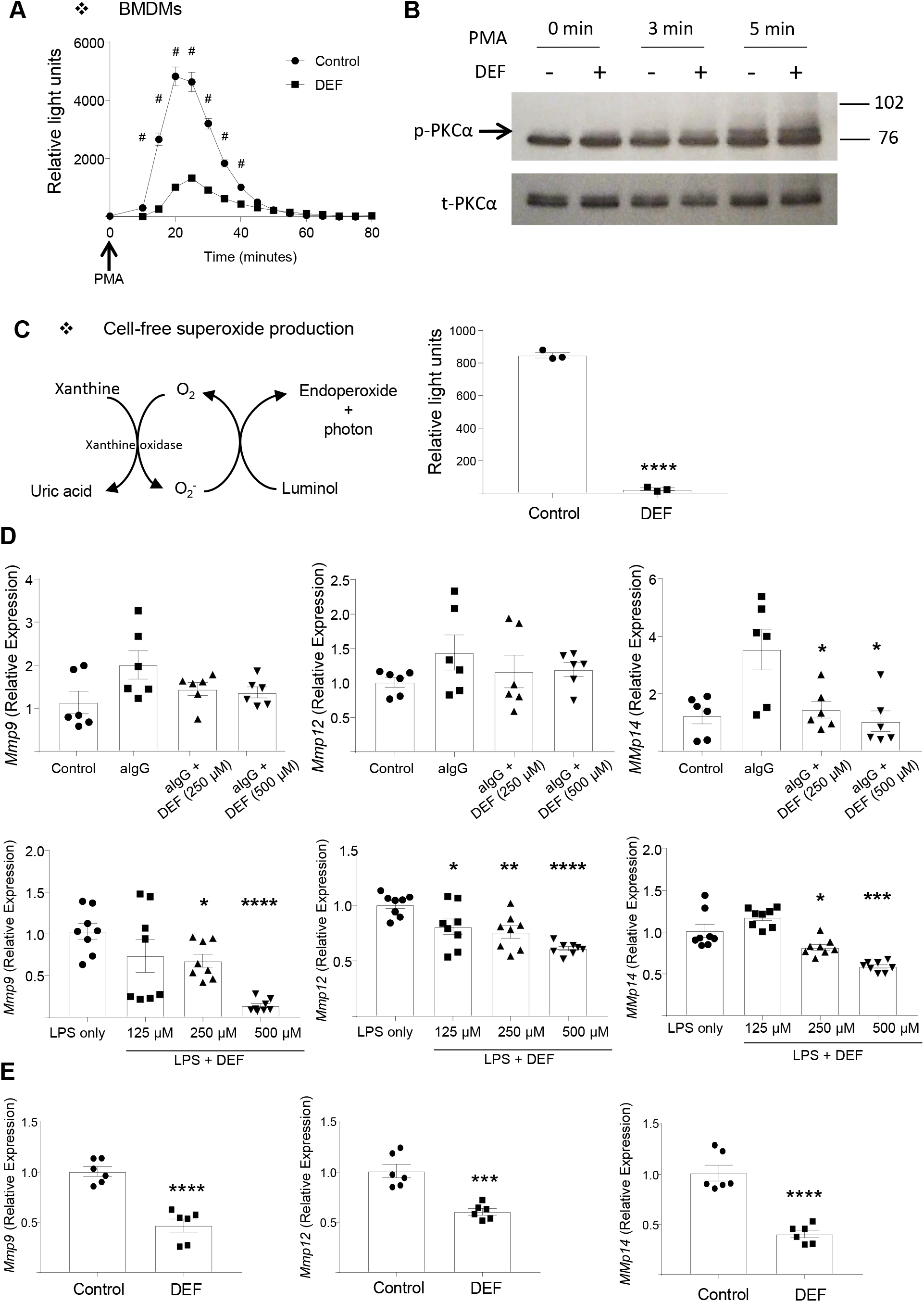
Deferiprone reduces superoxide anion production and down-regulates MMPs in macrophages (A) Superoxide chemiluminescence signal in BMDMs stimulated with 1 μM PMA with or without of deferiprone (DEF, 500 μM). (B) Western blot showing deferiprone does not prevent activation of PKCα in BMDMs induced by PMA stimulation. Note the time-dependent increase in PKCα phosphorylation (C) Superoxide chemiluminescence signal produced by xanthine-xanthine oxidase reaction in a cell free setup with or without deferiprone (DEF, 500 μM). (D) mRNA expression of *Mmp9, Mmp12* and *Mmp14* in BMDMs stimulated with heat aggregated IgG for 24h (aIgG, upper panel) or LPS for 3 hours (lower panel), with or without the indicated concentrations of DEF. (E) mRNA expression of *Mmp9, Mmp12* and *Mmp14* in nephritic glomeruli. Data are reported as mean±SEM. For all experiments, each group has at least 3 biological replicates with or without technical duplicates. \**P* ≤ 0.05; \*\**P* ≤ 0.01; \*\*\**P* ≤ 0.001; \*\*\*\**P* ≤ 0.0001; # *P* ≤ 0.0001.

Zinc-dependent matrix metalloproteinases (MMPs) are a family of proteinases involved in the degradation of extracellular matrix (ECM) components, and they have been implicated in the degradation of the glomerular basement membrane which associates with crescentic glomerulonephritis.^13,27,41–44^ The expression levels of MMP-9, −12, and −14 are under the regulation of superoxide anions,^26,45,46^ and in keeping with this, we found that deferiprone reduced *Mmp9, Mmp12,* and *Mmp14* mRNA levels in BMDMs activated by heat-aggregated IgG or LPS (Figure 3D). Furthermore, there was also a significant reduction of the above MMPs mRNA levels in the nephritic glomeruli of deferiprone-treated rats *in vivo,* when compared with the vehicle-treated rats. (Figure 3E).

### Deferiprone is Effective in Established Nephrotoxic Nephritis

To assess if deferiprone has therapeutic effects after the onset of glomerulonephritis, rats were treated with or without deferiprone from day 4 following the induction of nephrotoxic nephritis, which corresponds to the onset of renal damage and proteinuria. The rats were then sacrificed on day 10, at the peak of glomerular inflammation (Figure 4A).

**Figure 4.**
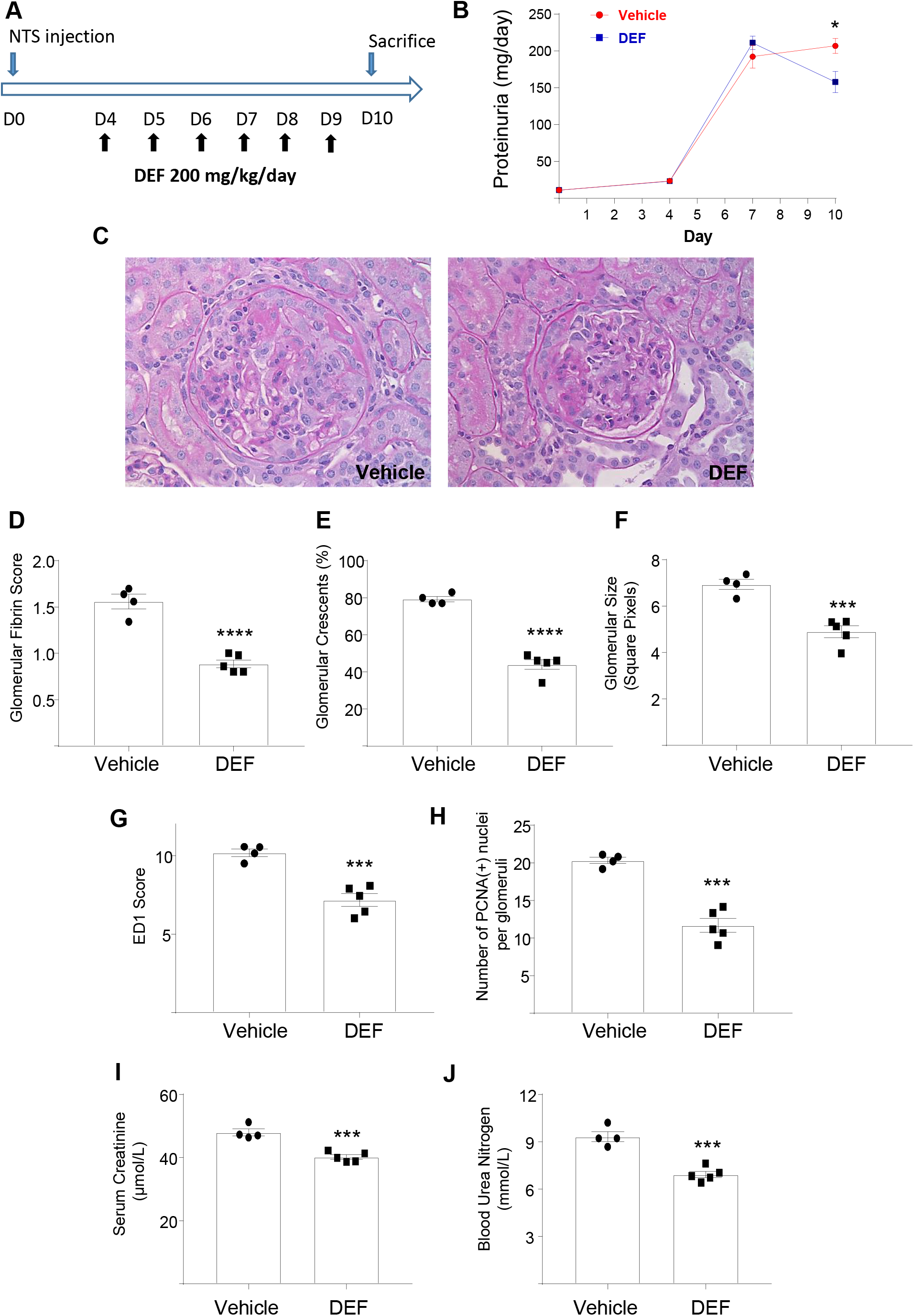
Deferiprone treatment attenuates established NTN; (A) Experimental design. (B) Amount of urinary protein levels at different time points during the course of NTN. (C) PAS staining showing representative glomerular morphology of the vehicle group (left) and the DEF group (right). (D) Glomerular fibrin score. (E) Percentage of glomeruli with crescents. (F) Glomerular size. (G) ED1 staining showing macrophages in glomeruli; ED1 score normalized to glomerular size. (H) PCNA staining showing proliferating cells in the glomerulus; number of nuclear PCNA(+) cells per glomerulus. (I) Serum creatinine (J) Blood urea nitrogen. Photographs original magnification, ×400. n=4 for the vehicle and n=5 for the deferiprone (DEF) groups. \**P* ≤ 0.05; \*\**P* ≤ 0.01; \*\*\**P* ≤ 0.001; \*\*\*\**P* ≤ 0.0001.

Baseline urinary protein levels were identical on day 0 before the injection of nephrotoxic serum. Proteinuria was detected on day 4 in both groups before the administration of vehicle or deferiprone. On day 7, both groups developed severe proteinuria. Notably, the progression of proteinuria was reversed in rats treated with deferiprone by day 10, as shown by lower levels when compared to either the treated group on day 7 or to their control counterparts on day 10 (Figure 4B).

The glomerular morphology of deferiprone-treated (DEF) rats on day 10 showed improved features when compared to vehicle-treated rats (Figure 4C). Fibrin deposition in glomeruli was significantly less in the deferiprone treated group (Figure 4D), and deferiprone also effectively reduced the formation of glomerular crescents (Figure 4E). The glomerular size was considerably smaller in the DEF group (Figure 4F), and monocyte/macrophage infiltration, as well as intraglomerular cell proliferation, were lowered by deferiprone (Figure 4G and H). Importantly, rats in the DEF group also showed better renal function with significantly lower levels of serum creatinine and blood urea nitrogen (Figure 4I and J). In summary, we propose that deferiprone inhibits cell proliferation and oxidative stress in macrophages through iron chelation and superoxide neutralising effects, respectively (Figure 5). The latter is associated with diminished levels of MCP-1 and MMPs, two important factors causing glomerular damage in the WKY NTN model.

**Figure 5.**
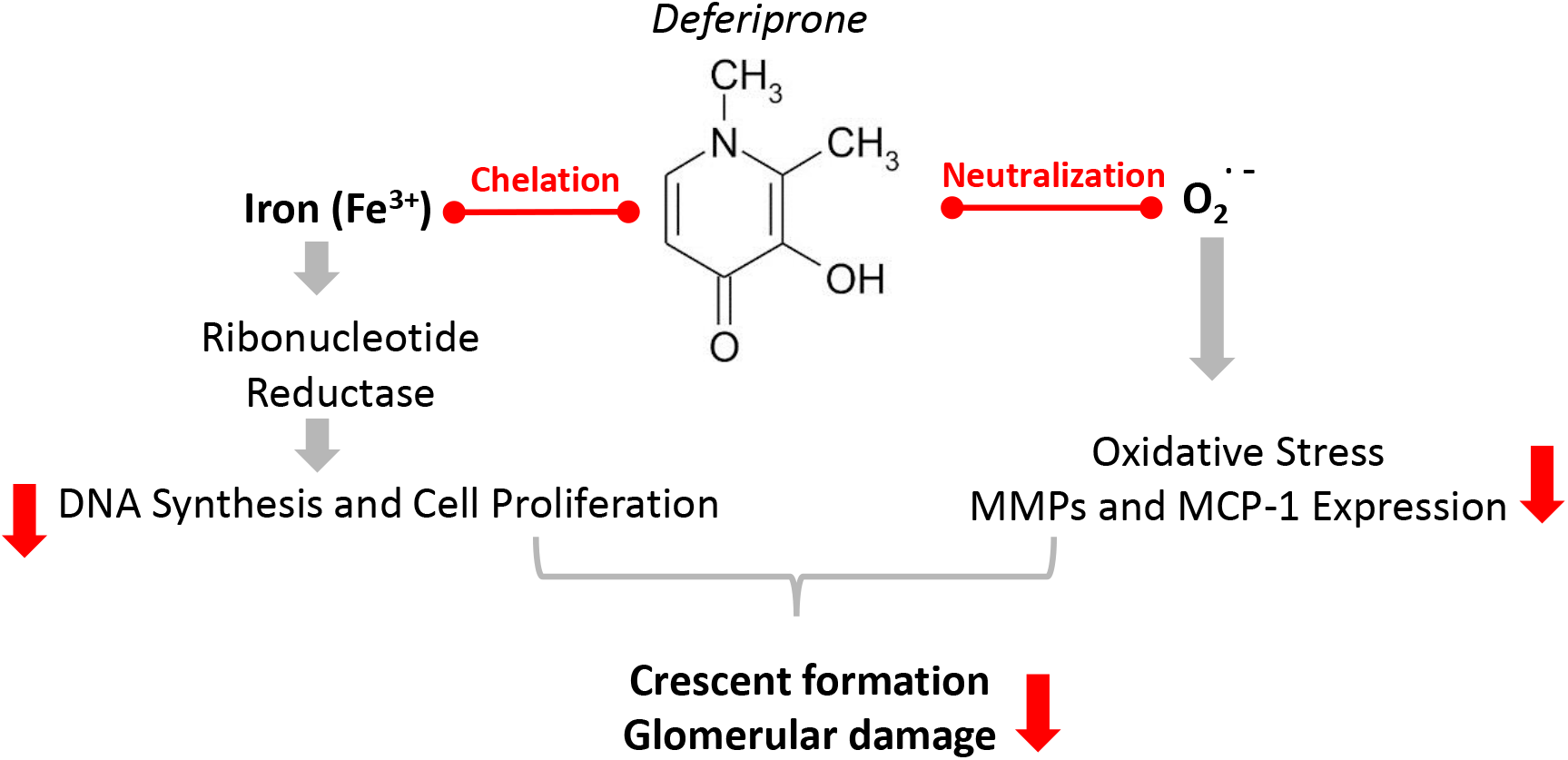
Schematic illustration of the proposed effects of deferiprone in macrophages. Deferiprone is likely to have renal protective effects through a combination of iron chelation and superoxide neutralization in macrophages, which could result in reduced expression of MMPs and MCP-1 as well as inhibition of cell proliferation, critical for the glomerular damage and crescent formation.

## Discussion

In this study we report the preventive and therapeutic effects of deferiprone, a widely used iron chelating drug, in an acute and severe rat model of crescentic glomerulonephritis. We show that the administration of deferiprone either before or during the course of experimental crescentic glomerulonephritis reduced disease severity assessed by morphological parameters (glomerular crescent formation and fibrin deposition) and functional indicators (proteinuria, serum creatinine, and blood urea nitrogen). Deferiprone’s effect can be attributed to its anti-oxidant and iron-chelating properties. Deferiprone markedly reduced superoxide anions produced by macrophages, possibly by directly neutralizing them, and therefore prevented subsequent oxidative damage and the expression of chemoattractant MCP-1 and MMPs. Deferiprone also prevented cell proliferation both *in vivo* and *in vitro,* possibly through its intracellular iron chelation effect.

Deferiprone (3-hydroxy-1,2-dimethylpyridin-4-one, also called L1) is an orally active iron chelator which preferentially binds to ferric iron (Fe^3+^) in a 3:1 configuration. It is an FDA-approved drug (known as FERRIPROX) used for the treatment of transfusion overload caused by thalassemia.^28^ When compared with other iron chelating drugs in clinical use, such as deferoxamine and deferasirox, preclinical and clinical toxicity evidence suggests that deferiprone can be used for long-term treatment of patients with non-iron loaded conditions, potentially extending its usage.^47^ Indeed, long-term deferiprone treatment has been used in trials involving non-iron loaded patients with Friedreich ataxia, neurodegenerative conditions, and diabetic nephropathy.^29^ As shown in the above studies and our current report, iron chelators such as deferiprone could have secondary therapeutic effects beyond the management of iron burden. Iron chelation prevents cell proliferation through inactivation of ribonucleotide reductase, one of the rate limiting enzymes in DNA synthesis.^18,48^ The antiproliferative effect of iron chelation is under investigation in anti-cancer therapies and tissue transplants.^19,49^

Superoxide production and the subsequent ROS formation, when not tightly controlled, elicit oxidative stress causing cell damage. Superoxide produced by macrophages is important for the development of crescentic glomerulonephritis,^36^ and ROS were reported to cause tissue damage in different models of experimental glomerulonephritis.^50^ In experimental anti-GBM and anti-Thy 1.1 glomerulonephritis, treatment with antioxidants attenuated proteinuria and glomerular injuries.^51–53^ Deferiprone has been shown as a potent antioxidant drug in many *in vitro* studies,^29,54–57^ as well as *in vivo* models,^58–60^ but not in experimental glomerulonephritis. It has recently been suggested that deferiprone inhibits the phosphorylation of PKCα, preventing the activation of its downstream target, NADPH oxidase, which is responsible for the production of superoxide anions.^40^ In addition, another study showed that deferiprone is effective at scavenging superoxide generated enzymatically *in vitro,* without the participation of PKCα or NADPH oxidase.^56^ Our results suggest that, in macrophages, the dominant antioxidant action of deferiprone is neutralization of superoxide anions, although we cannot exclude an inhibitory effect of deferiprone on NADPH oxidase.

In addition to their role in oxidative damage, superoxide anions are also responsible for MCP-1 expression in renal epithelial cells, vascular endothelial cells, and macrophages.^25,61,62^ The dose-dependent effect of deferiprone on *Ccl2* mRNA expression and MCP-1 synthesis in macrophages observed in our study can be attributed to deferiprone’s strong neutralizing effect on superoxide anions. Moreover, various MMPs including MMP-9, MMP-12, and MMP-14 are reported as tightly linked to glomerular damage in animal models of crescentic glomerulonephritis,^13,27,41,43,44^ as well as in human crescentic glomerulonephritis.^42^ Similarly, their expression are known to be regulated by superoxide,^26,45,46^ suggesting that oxidative stress is upstream of their expression and the subsequent glomerular inflammation. By neutralizing superoxide anions, deferiprone not only prevents tissue damage elicited by oxidative stress, but also reduces macrophage recruitment and the production of matrix degrading enzymes involved in glomerulonephritis. It has been shown that bone marrow derived cells account for 90% of the susceptibility to NTN in the WKY rat,^13,34^ supporting the beneficial effect of deferiprone on macrophage function. However, we cannot exclude that deferiprone treatment could also act on intrinsic renal cells.

The possibility of using deferiprone as a potential treatment for renal disease is attractive for several reasons. Currently there is no definitive treatment for crescentic glomerulonephritis, and general immunosuppressive medication such as steroids or cyclophosphamide cause severe side effects.

Deferiprone is an FDA-approved drug with world-wide usage in thalassemia patients for more than 15 years.^63^ When extending its long-term usage to non-iron loaded patients, such as those suffering from Friedreich ataxia and diabetic nephropathy, this did not result in any serious adverse events.^47^ Its oral administration further facilitates its potential long term usage in chronic kidney diseases. In keeping with this, deferiprone therapy achieved a significant reduction in proteinuria by 6 months in a small renal disease trial,^64^ strongly arguing in favour of its potential use in larger randomized trials.

In conclusion, our study showed preventive and therapeutic effects of deferiprone in an acute and severe model of crescentic glomerulonephritis in the rat. Our *in vitro* experiments in macrophages showed that deferiprone chelated iron, inhibited cell proliferation, and neutralized superoxide anions. *In vivo*, deferiprone treatment was associated with reduced inflammatory cell recruitment and their further release of MMPs. Combined with the recent findings highlighting the importance of iron in immune cell proliferation, differentiation, and function, further investigation of mechanisms behind macrophage iron regulation and oxidative stress in cells will offer a new therapeutic angle in macrophage-dependent kidney diseases. Our current study provides strong evidence that deferiprone could be a novel therapy in the management of crescentic glomerulonephritis.

## Concise Methods

### Nephrotoxic Nephritis and Treatment with Deferiprone

WKY rats (Charles River UK) weighing 300-360 g were used for the induction of NTN. Nephrotoxic serum was prepared in rabbits and NTN was induced in male WKY rats by intravenous injection as previously described.^8^ Deferiprone (3-Hydroxy-1,2-dimethyl-4(1H)-pyridone) was purchased from Sigma-Aldrich. Groups of animal were given either deferiprone (200 mg/kg/day in 0.1% carboxymethyl cellulose as vehicle; divided in two doses by oral gavage) or vehicle only as control. Rats were individually housed in metabolic cages overnight for urine collection, and urinary protein levels were determined by sulphosalicylic acid method.^65^ Serum blood urea nitrogen and creatinine were measured using an Abbott Architect c8000 clinical chemistry analyser. Animal study procedures were performed in accordance with the United Kingdom Animals (Scientific Procedures Act).

### Light Microscopy and Immunohistochemistry

Paraffin-embedded kidneys were sectioned for periodic acid-Schiff (PAS) staining. Fibrin deposition was scored as the number of glomerular quadrants involved from ‘0’ (absence of fibrin) to ‘4+’ (fibrin occupying more than ¾ of the whole glomerulus) in 50 consecutive glomeruli. Crescent formation was recorded as presence / absence in 100 consecutive glomeruli and presented as percentage of glomeruli with crescents in the kidney from each rat. Immunohistochemistry was performed and developed with EnVision+ System-HRP (K4007, Dako) using mouse anti-rat ED1 (MCA341R, BioRad) and anti-PCNA [PC10] (ab29, abcam) antibodies. Pictures of 20 consecutive glomeruli from each ED1 stained slide were taken with a QImaging Retiga 2000R Scientific CCD Camera, using Image-Pro^®^ Plus Version 7.0 software. The size of the glomeruli and ED1 stained area in glomeruli were measured using ImageJ software. The number of anti-PCNA stained cells in 20 consecutive glomeruli were counted. All histological and immunohistochemical analyses were performed in a blinded manner.

### Bone Marrow Derived Macrophage Culture and Stimulation

Femurs were obtained from adult WKY rats and flushed with Hank’s Balanced Salt solution (Gibco). BMDMs were prepared by incubating bone marrow-derived total cells in either L929 cell-conditioned media or culture media containing M-CSF. For heat-aggregated IgG experiments, BMDMs were changed to serum-free medium 24 hours before stimulation and incubated with serum-free medium containing either deferiprone or control medium for 2 hours before addition of heat-aggregated rat IgG (Sigma). Following 24 hours, supernatants were collected for measurement of MCP-1, and RNA were collected. For LPS stimulation, BMDMs were incubated with deferiprone during differentiation for 36 hours, treated with LPS (100 ng/mL) for 3 hours, and RNA was collected. The cell viability was assessed by Trypan blue stain (Invitrogen).

### Quantitative RT-PCR

Total RNA was extracted from BMDMs or nephritic glomeruli using the TRIzol reagent (Invitrogen) according to the manufacturer’s instructions, and cDNA was synthesized using iScript™ cDNA Synthesis Kit (Bio-Rad). A total of 10 ng cDNA for each sample was used. All quantitative RT-PCRs were performed on a ViaA 7 Real-Time PCR System (Life Technologies) using Brilliant II SYBR Green QPCR Master Mix (Agilent), followed by ViiA 7 RUO Software for the determination of Ct values. Results were analysed using the comparative Ct method, and each sample was normalized to the reference mRNA level of *Hprt* gene, to account for any potential cDNA loading differences. The primer sequences are available upon request.

### Western Blotting

BMDMs were homogenized and lysed in 1% NP-40 lysis buffer with protease and phosphatase inhibitors, mixed with 2X Laemmli sample buffer (Bio-Rad), resolved by SDS-PAGE, transferred to polyvinylidene difluoride membranes, and subjected to immunoblotting with either rabbit monoclonal anti-PKCα [phospho S657] (ab180848, abcam) or anti-PKCα [Y124] (ab32376, abcam). Following incubation with secondary antibodies, the probed proteins were detected using SuperSignal West Femto Chemiluminescent Substrate (Thermo Fisher Scientific).

### Measurement of Superoxide Anion Production

Superoxide anions were assessed by chemiluminescence assays (LumiMax Superoxide Anion Detection Kit, Agilent; Superoxide Anion Assay Kit, Sigma), wherein relative luminescence unit (RLU) values corresponded to superoxide levels produced by BMDMs, as previously described with minor modifications.^37^ Briefly, day-5 BMDMs were re-suspended in cell dissociation solution (Sigma) and re-plated in a 96-well optical flat bottom plate (CELLSTAR^®^) at a cell density of 2.5×10^5^ cells per well, allowed to adhere and cultured in DMEM medium with 10% FBS overnight. The experiment was performed with four biological and two technical replicates. Two hours prior to the assay, deferiprone and vehicle (sterile water) were added into experimental group and control group wells, respectively, to achieve a final concentration of 500 μM deferiprone during the assay. PMA (1 μM, Sigma) was used to generate superoxide production by BMDMs. Chemiluminescence was detected using a FLUOstar Galaxy plate reader (BMG LABTECH), and the RLU values were detected for a total period of 80 minutes. The cell-free superoxide production and the neutralizing effect of deferiprone was measured by xanthine/xanthine oxidase reaction in the same chemiluminescence system (Superoxide Anion Assay Kit, Sigma).

### Statistics

Statistical analysis was conducted using GraphPad Prism 7.0 (GraphPad Software Inc.). All data were reported as mean with standard error of mean (SEM) unless otherwise stated. Comparison between groups was done by two-tailed Student’s t test.

## Acknowledgments

This work was supported by the Medical Research Council (MR/M004716/1 and MR/N01121X/1 to J.B.) and by Kidney Research UK (RP9/2013 to J.B.).

